# Integrating EM and Patch-seq data: Synaptic connectivity and target specificity of predicted Sst transcriptomic types

**DOI:** 10.1101/2023.03.22.533857

**Authors:** C.R. Gamlin, C.M. Schneider-Mizell, M. Mallory, L. Elabbady, N. Gouwens, G. Williams, A. Mukora, R. Dalley, A. Bodor, D. Brittain, J. Buchanan, D. Bumbarger, D. Kapner, S. Kinn, G. Mahalingam, S. Seshamani, M. Takeno, R. Torres, W. Yin, P.R. Nicovich, J.A. Bae, M.A. Castro, S. Dorkenwald, A. Halageri, Z. Jia, C. Jordan, N. Kemnitz, K. Lee, K. Li, R. Lu, T. Macrina, E. Mitchell, S.S. Mondal, S. Mu, B. Nehoran, S. Popovych, W. Silversmith, N.L. Turner, W. Wong, J. Wu, S. Yu, J. Berg, T. Jarsky, B. Lee, H.S. Seung, H. Zeng, R.C. Reid, F. Collman, N.M. da Costa, S. A. Sorensen

**Author notes:** denotes corresponding author.

## Abstract

Neural circuit function is shaped both by the cell types that comprise the circuit and the connections between those cell types^1^. Neural cell types have previously been defined by morphology ^2, 3^, electrophysiology ^4, 5^, transcriptomic expression ^6–8^, connectivity ^9–13^, or even a combination of such modalities ^14–16^. More recently, the Patch-seq technique has enabled the characterization of morphology (M), electrophysiology (E), and transcriptomic (T) properties from individual cells ^17–20^. Using this technique, these properties were integrated to define 28, inhibitory multimodal, MET-types in mouse primary visual cortex ^21^. It is unknown how these MET-types connect within the broader cortical circuitry however. Here we show that we can predict the MET-type identity of inhibitory cells within a large-scale electron microscopy (EM) dataset and these MET-types have distinct ultrastructural features and synapse connectivity patterns. We found that EM Martinotti cells, a well defined morphological cell type ^22, 23^ known to be Somatostatin positive (Sst+) ^24, 25^, were successfully predicted to belong to Sst+ MET-types. Each identified MET-type had distinct axon myelination patterns and synapsed onto specific excitatory targets. Our results demonstrate that morphological features can be used to link cell type identities across imaging modalities, which enables further comparison of connectivity in relation to transcriptomic or electrophysiological properties. Furthermore, our results show that MET-types have distinct connectivity patterns, supporting the use of MET-types and connectivity to meaningfully define cell types.

## Introduction

To understand the function of a complex biological structure, such as the brain, we must first define the building blocks, or cell types, that make up the structure. Next, we must determine how those building blocks fit together. In mouse primary visual cortex (VISp), at least 28 inhibitory cell types have been defined by their concordant morphology (M), electrophysiology (E), and transcriptomic expression (T) (MET-types)^21^ from data obtained from the Patch-seq method ^26^. These MET-types align well with previously characterized inhibitory cell types (e.g. somatostatin, Parvalbumin) ^23, 27, 28^. Using these MET-types as building blocks, we aim to determine the synaptic connectivity of these identifiable inhibitory cell types.

The Patch-seq method enables a single cell to be recorded and filled with dye, and then the cell’s contents are extracted for subsequent single-cell RNA-sequencing using the whole-cell patch-clamp technique ^17–19, 29^. The cell’s transcriptomic expression profile is determined from the sequenced cellular material and is mapped onto existing cell taxonomies to assign a transcriptomic type (t-type) ^7, 8^. Some MET-types contain a single t-type, and some t-types are divided across MET-types, but most commonly inhibitory MET-types contain several neighboring t-types in the taxonomy that also share similar morphological and electrophysiological properties. We leverage the known morphological features of inhibitory MET-types as defined by Patch-seq to predict the MET-type identity of cells in a large-scale electron microscopy (EM) dataset from mouse VISp (collected as part of the MICrONs project) ^30^. We focused on Martinotti cells (MCs) in layers 4 and 5 (L4, L5), which have distinct morphological features ^23, 31, 32^, are known to be somatostatin positive (*Sst+*)^24, 25^, and span six Sst+ MET-types^21^. Additionally, previous functional studies suggest that MCs may connect broadly in a ‘blanket of inhibition’ ^33–35^ or inhibit a subset of excitatory targets within and across cortical layers ^32, 36–39^. Finally, in a concurrent study ^40^ individual MCs are found to preferentially target different excitatory cell types, however, it remains unknown whether Sst-MET-types have distinct synaptic connectivity profiles especially as morphologically similar neurons can have distinct connectivity profiles ^41^.

We reconstructed Martinotti cells from the EM dataset and predicted their MET-type identity using a random forest classifier trained on morphological features from Patch-seq cells. We find that each predicted MET-type differs in connectivity patterns with respect to both target cell subclasses and the number of synapses onto individual postsynaptic targets (multi-synaptic connections). We see biased connectivity onto excitatory targets both across and within cortical layers. We also find that MET-types can differ in total number and average size of output synapses as well as axonal myelination patterns. These differences in postsynaptic connectivity and myelination likely support distinct functional roles for inhibitory MET-types. Overall, by linking Patch-seq and EM data through neuron morphology, we have developed a method by which we can interrogate the relationships between transcriptomically-defined cell types, morphology, electrophysiology, and synaptic connectivity.

## Results

### Morphological features from inhibitory EM reconstructions are comparable to features from Patch-seq data

We identified and reconstructed 16 MCs with somas in L4 and L5 from a large-scale serial-section EM dataset (Fig 1A)^30^. Martinotti cells were identified by their sparsely spiny dendrites, an axon emerging from the pia-side of the soma, and a primary axon branch that reached L1^23^. We aligned the reconstructions to an average cortical layer space using a pipeline developed for Patch-seq data and calculated the morphological features originally used to characterize the morphology of neurons from *in vitro* slices of tissue ^16, 21^. These features include measurements such as total axon length, maximum path distance, and total number of branches for both axon and dendrites from skeletonized representations. Here we compare morphological features from cells reconstructed in the EM and Patch-seq pipelines (Supplemental Fig 1, 2) to evaluate feature alignment between the datasets.

**Fig 1:**
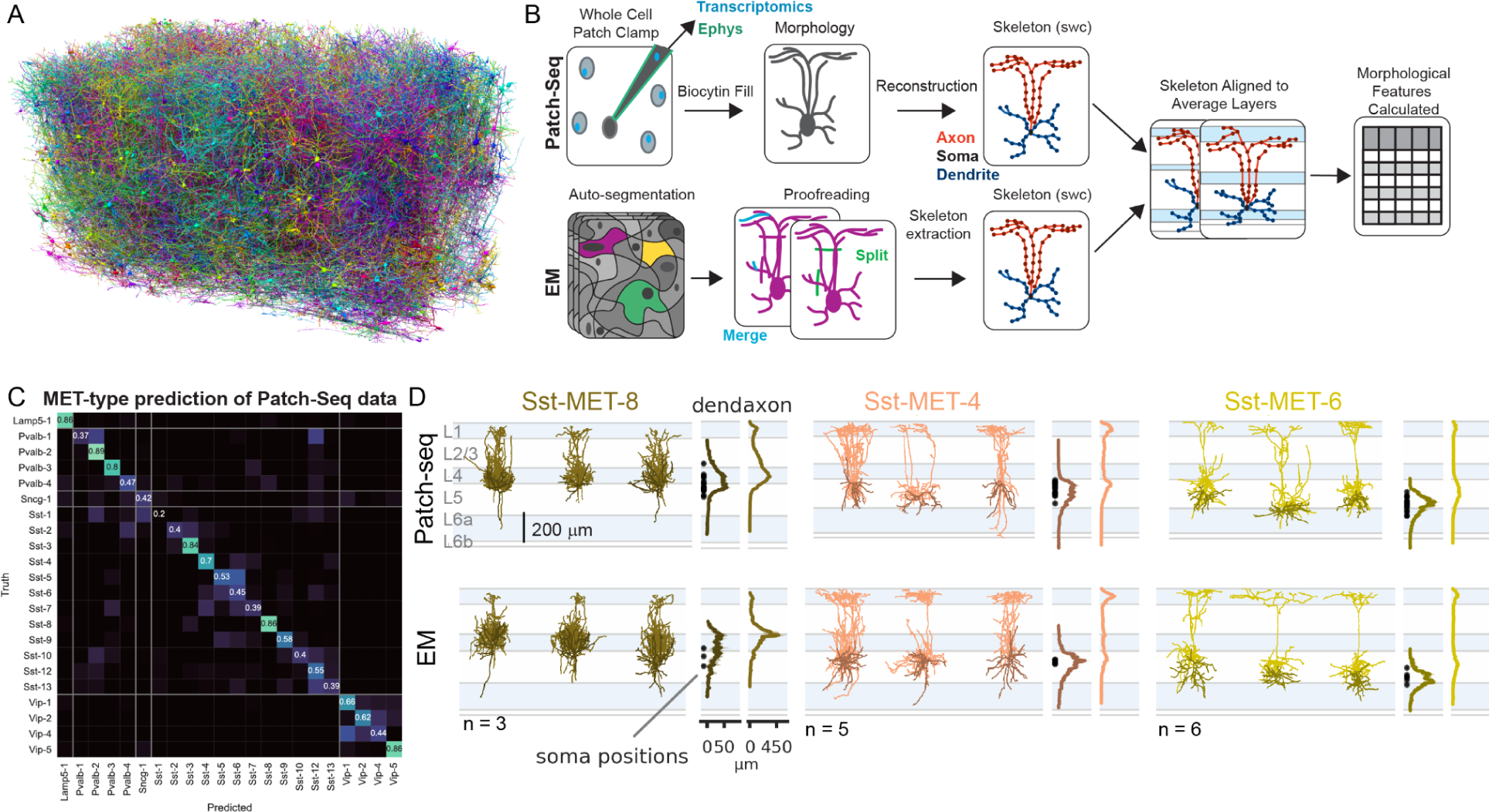
Comparison of EM and Patch-seq pipelines and morphologies. A. A subset of morphologies available from large-scale mm3 EM volume. Each cell is in a different color. B. Schematic representing the Patch-seq and EM pipelines for generating morphological reconstructions and comparison of features across pipelines. C. Confusion matrix of random forest classifier (RFC) MET-type predictions. T he RFC was trained on morphological features of inhibitory Patch-seq data. We used this classifier to predict MET-type identity of EM cells. D. (Top) Example Patch-Seq morphology and average axon/dendrite histograms of MET-types. (Bottom) EM cells morphology and average axon/dendrite histograms grouped by predicted MET-type (MET-8 n = 3; MET-4 n=5, MET-6 n = 6).

For all morphological features calculated (43/43), we found that the values from EM MCs fall within the range of values calculated from all inhibitory Patch-seq cells (which include cells from the *Sst*, *Pvalb*, Lamp5, Sncg, and *Vip* subclasses) for that specific feature, despite the different sampling methods used to generate each dataset (Supplemental Fig 1). We also found that all of the features of our EM MC cells (presumed to be from the Sst subclass) fall within the range of values calculated from only the *Sst*+ Patch-seq cells (Supplemental Fig 1, 2). We therefore proceeded to use these features to predict MET-type identity for EM MCs.

### Martinotti cells from EM are predicted to belong to *Sst*+ MET-types

Using a random forest classifier trained on morphological features of all inhibitory Patch-seq cells (Fig 1C), we predicted the MET-type for each EM MC cell (Fig 1D). Every reconstructed EM MC cell was predicted to belong to an *Sst*+ MET-type with high probability except for one (see Supplemental Table 1 for MET-type prediction probability), supporting the use of these morphological features across datasets to predict MET-type identity.

The MET-types^21^ represent cell types with concordant morphology, electrophysiology, and transcriptome. Some of these MET-types consist of cells that map to a single transcriptomic type (e.g. Sst Chodl and Sst Hpse Cbln4) whereas other MET-types contain cells that map to several, transcriptomically similar t-types, typically from neighboring branches of the transcriptomic dendrogram ^8^. The EM MC MET-type predictions included 5 *Sst+* MET-types described below.

Three EM cells are predicted to belong to the Sst-MET-8 type. Sst-MET-8 consists of cells from a single t-type, Hpse Cbln4, which have somas located in L4 and upper L5. Hpse Cbln4 cells are described as L5 non-Martinotti cells in somatosensory cortex ^29, 38^, but are observed to have some L1 projection in primary visual cortex and hence are considered Martinotti cells ^21, 29^.

Five EM cells are predicted to be Sst-MET-4. The Sst-MET-4 type consists mainly of cells from the Sst Calb2 Pdlim5 t-type. Interestingly, the Sst Calb2 Pdlim5 t-type is split across layers with one group found in L2/3 (Sst-MET-3) and another in upper L5 (Sst-MET-4) due to differences in their soma location, laminar innervation pattern, and overall size^21^, but all Sst Calb2 Pdlim5 cells are characterized by a L1-dominant axon lamination (‘Martinotti’) pattern ^23, 42^. Sst-MET-4 cells have previously been compared to L5 fanning-out Martinotti cells ^32, 37^.

Six EM cells are predicted to be Sst-MET-6. Sst-MET-6 cells were previously described as having “T-shaped” morphology with an axon branch that reaches L1 but has more dominant L5 innervation^37^. The Sst-MET-6 type is composed of several t-types that are in proximity along the transcriptomic dendrogram including Sst Myh8 Etv1 and Sst Chrna2 Glra3. *Chrna2*+ Martinotti cells have previously been described^36^ as preferentially connecting to “type A” thick tufted pyramidal neurons over thin tufted “type B” neurons in layer 5.

Lastly, one cell is predicted to be in the Sst-MET-5 type and one is predicted to be Sst-MET-9. Sst-MET-5 was previously described as morphologically similar to Sst-MET-6^21^. The Sst-MET-5 type is comprised of both Myh8 Etv1 and Nr2f2 Necab1 cells (similar to fanning-out Martinotti cells)^21, 32, 37^. The Sst-MET-9 type is primarily made up of the Tac2 Tacstd t-type and has an axon that reaches layer 1 but has a peak in layer 5.

These results demonstrate that we can reliably assign a morphological, electrophysiological, and transcriptomic identity to neurons characterized in an EM volume using local dendritic and axonal morphology. In the remainder of the paper, we will refer to these “predicted MET-types’’ as MET-types and focus our analysis on those with at least 3 cells (MET-8, 4, 6).

### Output synapse number and axonal myelination patterns vary by MET-type

#### I. Output synapse features

In the EM volume, most of output synapses (average 78% of all output synapses, Supplemental Table 2) along the reconstructed axons can be mapped onto a single postsynaptic target with a predicted cell subclass–based on somatic features^43^. This high rate of mapping onto target cells allowed us to confidently investigate the relationship between MET-type identity and connectivity statistics of Sst-MET-types 8, 4, and 6 (Fig 2A-B).

**Fig 2:**
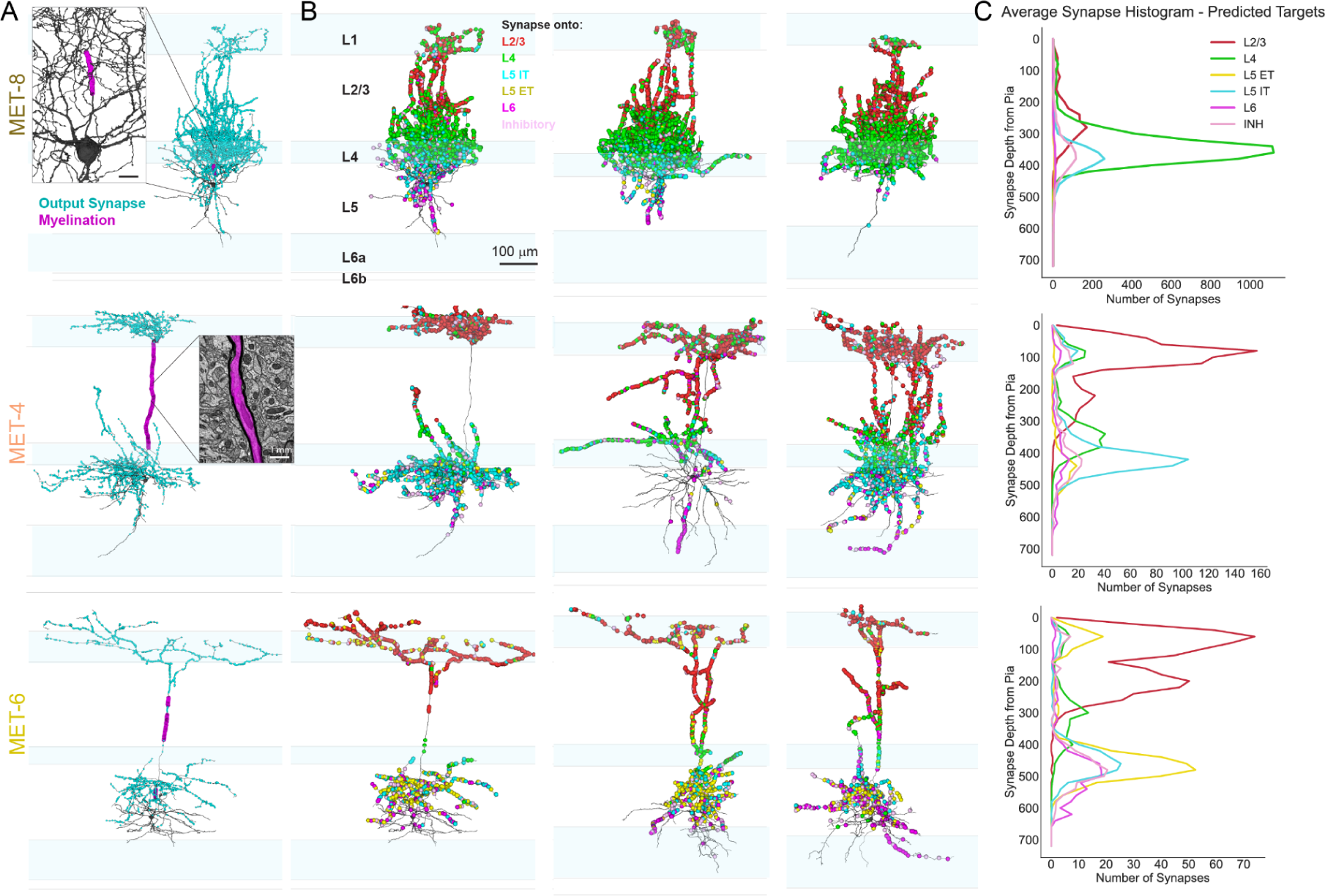
Output synapses and myelination of MET-types. A. Example cell from each predicted MET-type showing output synapses (cyan dots) and myelination (magenta). Insets show myelination close to soma of predicted MET-8 cell and myelinated axon in EM. B. Same exmple cells (plus additional examples) from each predicted MET-type with output synapses color-coded by synapse target (as identified by a classifier trained on somatic features - Elabbady et al.,^43^). C. Average histogram of synapses onto targets by predicted MET-type.

We find that MET-8 cells form significantly more output synapses (9046 ± 1336, mean ± SEM) than either the MET-4 (2181 ± 297) or MET-6 cells (1510 ± 161) (Fig 2A,C; for quantification see Supplemental Fig 3A,C). Additional synaptic features are described in Supplemental Fig 3. We also find significant differences in output synapse size across MET-types, discussed in greater detail in the Single vs. Multi-synaptic connections section below.

#### II. Myelination patterns

EM data includes information about the ultrastructure of processes such as myelination along the axon, which can influence the biophysics of a cell ^44^. While inhibitory neurons ^45, 46^, including Martinotti cells^47^, have previously been shown to be myelinated, to date little is known regarding the relationship between myelination pattern and Sst cell types and biophysical properties. We identified potential regions of myelinated axon by the absence of output synapses and manually annotated the start and end points of myelinated axonal segments of these neurons from the EM images. We compared the number and length of myelinated segments across MET-types (Fig 2A, Supplemental Fig 3B).

We found distinct myelination patterns across the three MET-types (Fig 2A, Supplemental Figure 4). MET-4 cells had a major ascending axon collateral with myelination along its length (through L4 and L2/3). MET-8 cells on the other hand were mostly myelinated along a short stretch of the primary axon branch located near the soma, which rarely extended into layer 2/3. In contrast, MET-6 cells were either not myelinated or sparsely myelinated with a less clear pattern (Fig 2A, Supplemental Figure 3B, 4). In total we find that most of the reconstructed cells had some portion of their axon myelinated, but the number of myelinated segments and total path length of the myelination varied by MET-type (Supplemental Figure 3B). All MET-8 (3/3) and MET-4 (5/5), as well as ∼83% (5/6) of MET-6 cells have some myelination. The MET-5 (1/1) and MET-9 (1/1) cells also have myelin.

The ascending axon stalk of MET-4 cells is fully myelinated by multiple segments (∼8) separated by nodes of Ranvier. Consequently we find that they have approximately three-times as many segments as MET-8 (∼3) and five-times as many segments as MET-6 (∼2). There is no significant difference in the length of individual myelin segments. MET-4 cells have approximately 260 µm of total length of myelination, which is nearly four-times the length of myelin of MET-8 (∼69 µm) and seven-times the length of myelin of MET-6 (36 µm) (Supplemental Fig 3B).

### Synaptic targets and connectivity patterns vary by MET-type

#### I. Synaptic connectivity

Excitatory cortical cell subclasses are named for both the layer in which they reside (i.e. L2/3) and for where they project their axons. Intra-telencephalic (IT) neurons project within the cortex and extra-telencephalic (ET) neurons project beyond the cortex^48^. IT and ET (sometimes called PT) neurons are functionally distinct and differentially implicated in several diseases^49^. We determined the synaptic connectivity pattern of each Sst+ MET-type using automated methods to detect synapses^50^ and assign target subclass identity^43^. We also manually confirmed the presence of a synapse and target identity for 5% of the synapses for a subset of cells (see Methods for additional details). Here we report the number of output synapses formed onto predicted cell subclasses (Fig 2B-C). Since MET-types vary in total number of output synapses, to compare connectivity patterns, we measured both total number and fraction of synapses from individual cells that targeted each predicted post-synaptic cell subclass (Supplemental Fig 3C).

MET-8 cells preferentially target L4 IT pyramidal cells (62.3% ± 3.8, mean± SEM, same below). The next major targets are L5 IT and L2/3 PC (pyramidal cells) (13.2% ± 1.5 and 13.8% ± 4.0, respectively). This is largely consistent with the laminar innervation pattern of its axon, though layer 4 contains many apical dendrites which are not from layer 4 neurons.

MET-4 cells predominantly target L2/3 (37.7% ± 4.2) followed by L5 IT cells (25.7% ± 7.1). L4 ITs and inhibitory neurons receive fewer synapses (13.3% ± 4.7 and 11.9% ± 1.0, respectively). One MET-4 cell does preferentially synapse with L5 ETs rather than L5 IT, which may indicate variability either in targeting for this MET-type or misassignment of MET-type for this cell. Further characterization of the connectivity of other MET-types will help elucidate their intrinsic variability.

MET-6 cells form most synapses onto L2/3 pyramidal cells (35.6% ± 1.7%) followed by L5 ET and L6 pyramidal cells (22.3% ± 4.0 and 13.8 ± 3.5%, respectively). MET-6 cells target L5 IT and inhibitory cells to a lesser extent (L5 IT: 11.4% ± 0.7 and inhibitory: 11.1% ± 0.1). Comparing the connectivity of MET-4 and MET-6 cells is of particular interest, because they both have significant axon projections in layer 1 and layer 5, and some cells from both groups could both be considered “T-shaped” MCs^37^, yet are predicted to belong to distinct Sst+ MET-types and have distinct connectivity profiles.

Despite having somas in layer 5, we find that the major targets for both Sst-MET-4 and MET-6 are L2/3 pyramidal cells. This contrasts with the view that the primary role of L5 MCs is to inhibit the apical tufts of layer 5 excitatory neurons ^51, 52^ though these synapses are also present. As described above, each MET-type preferentially synapses with a specific layer 5 excitatory cell subclass. We find that the synapses in layer 1 formed onto the apical tufts of layer 5 targets are onto each type’s preferred target subclass (Figure 2C). While MC axons have previously been described as overlapping with the basal dendrites of their L5 targets ^23, 32, 53–55^, here we show that more synapses onto the preferred L5 targets are formed within L5 than L1 (see Fig 2C: peaks at 0-100 µm from pia) as opposed to purely targeting apical tufts.

#### II. Target cell locations

The observation that Sst-MET types have distinct output synapse distributions suggests that Sst+ cells are not indiscriminately synapsing onto all subclasses (i.e. ET vs IT), but it does not directly measure whether all available cells within a subclass are being innervated, as predicted by a “blanket” inhibition model^33^. Therefore, we calculated the percentage of cells of each target subclass within a given radial distance that receive synapses from the presynaptic MET-type. We find that connectivity rates peak within 100 µm for every subclass however, the percentages vary by MET-type and target cell subclass and few are near 100% (Fig 3A).

**Fig 3:**
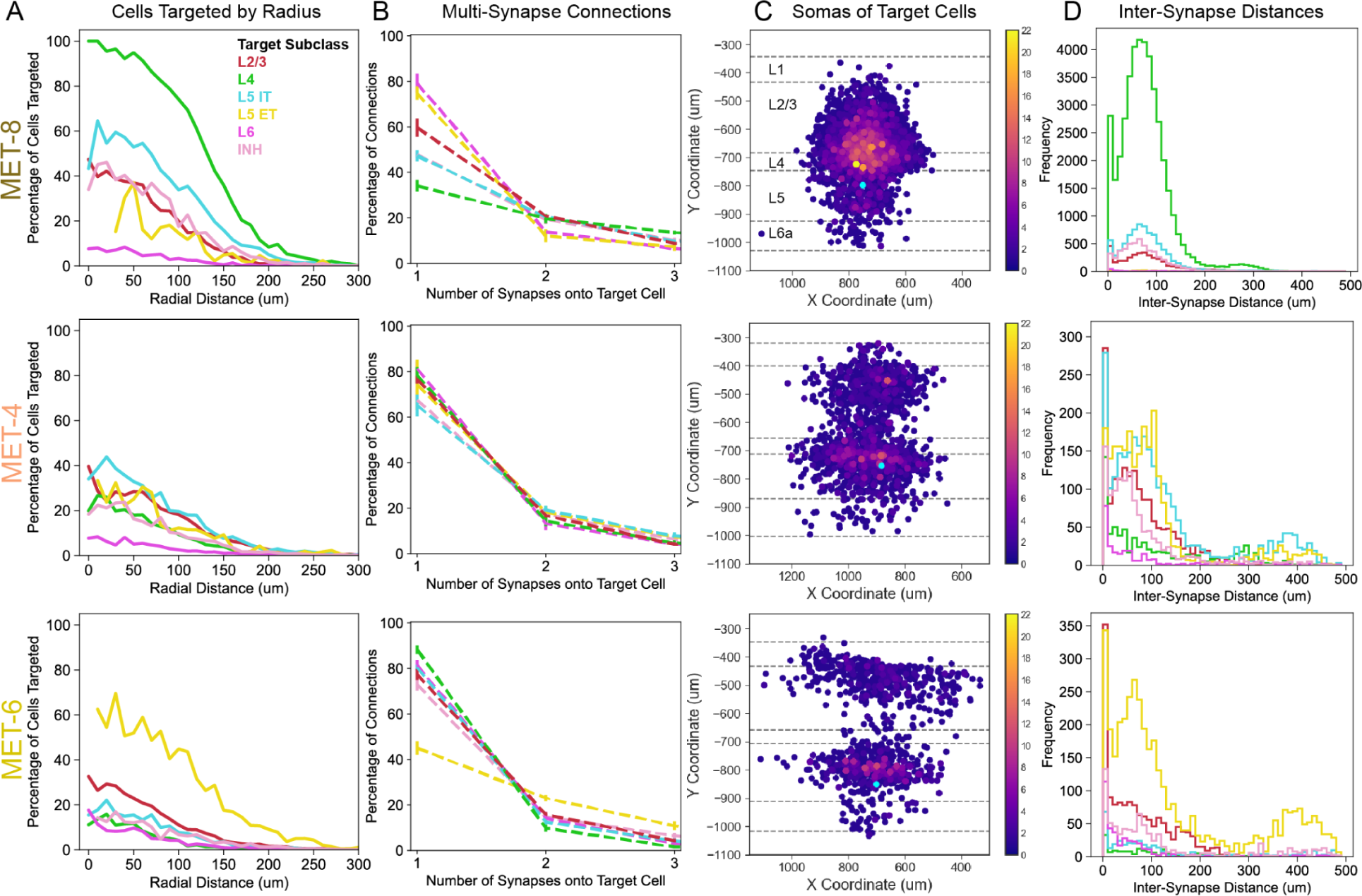
Exploring pair-wise connectivity of MET-types and postsynaptic targets. A. Average histogram of percentage of cells of each cell type targeted as a function of radial (xz) distance between presynaptic EM cell and postsynaptic target. B. Percentage of connections that contain a single vs 2 or 3 synapses for each target cell type across predicted MET-types. (e.g. Predicted MET-6 cells form more multi-synaptic contacts onto L5 ET targets vs other cell types).C. Example cell from each predicted MET-type showing soma locations of postsynaptic targets. Somas are color-coded to indicate the number of synapses that cell receives from the presynaptic cell (soma in cyan). All examples use the same scale. D. Histograms of inter-synaptic distances onto targets types (distances calculated per postsynaptic target).

MET-8, MET-4 and MET-6 cells target ∼40% or fewer of the available cells from each subclass within 50 µm, with a few notable exceptions, indicating that these inhibitory cells are not indiscriminately synapsing with every available neuron, or “blanketing”, all surrounding neurons with inhibition. However, MET-8 cells target nearly 100% of all L4 cells within 50 µm of the soma and 80% within 100 µm of the soma, producing a “blanket of inhibition” for those targets. Thus, a “blanket of inhibition” can be seen when analyzing specific pre- and post-synaptic cell subclass. Like the MET-8 cells, MET-6 cells target a high percentage of one of their preferred postsynaptic targets (nearly 70% of all L5 ET neurons within 100 µm), but not others (only approximately 30% of the available L2/3 cells within 50 µm radial distance). MET-4 cells do not target a large percentage of the available cells of any subclass (less than 40% of all available cells across subclasses within 50 µm), not even their preferred targets as assessed by percentage of output synapses (L2/3 and L5 IT cells). Thus, each MET-type employs a distinct inhibition pattern – inhibiting varying fractions of target subclasses in line with preferences (calculated from percentage of synapses) reaching nearly 100% of the preferred subclass (MET-8); inhibiting a similarly small fraction of all target subclasses (MET-4); or inhibiting a small fraction of target subclasses except for the most preferred “local” (L5 ET) target (MET-6). Our findings align well with paired recordings showing high connection probability from L4 Sst cells to L4 pyramidal cells and L5 Sst cells onto L5 ET versus IT cells ^56^. Our radial distance measurements also align with previous studies in brain slices showing most connections from Sst cells onto excitatory targets occur within 200 µm lateral distance from the cell soma ^56, 57^.

#### III. Single vs Multi-synapse connections

Given that individual neurons do not connect to every surrounding neuron, the differences in the output distribution of synapses between the three MET types may be due to differences in the probability of connection to targets (Fig 3A), but also could be due to shifts in the average number of synapses per subclass connection (Fig 3B). MET-types can form single or multiple synapses onto individual postsynaptic targets. We calculated the average number of synapses onto each target subclass for each MET-type (Fig 3B). We found that most connections, including those onto L2/3 pyramidal cells, contain a single synapse. However, the relative fraction of single synapse and multi-synapse connections was strongly modulated by the postsynaptic target subclass (Fig 3B).

Though most of the Sst-MET-type connections contain single-synapses, there are a few notable exceptions. MET-8 cells make the most multi-synaptic contacts onto their major targets: L4, L5 IT, and inhibitory cells. Specifically, greater than 60% of all connections onto L4 cells are multi-synaptic (more than 1 synapse) and greater than 50% of connections onto L5 IT and inhibitory targets are multi-synaptic. Some targets receive greater than 20 synapses from a single MET-8 cell. MET-4 cells form the fewest multi-synaptic connections and show the least modulation with respect to target cell subclass. MET-6 also forms few multi-synaptic connections overall, including onto their major target (L2/3 PCs), but instead form highly multi-synaptic connections onto L5 ET pyramidal cells (∼50% of connections are multi-synaptic). In summary, MET-8 forms more multi-synaptic connections onto its major postsynaptic targets, MET-4 does not form many multi-synaptic connections, and MET-6 forms multi-synaptic connections onto one specific target subclass. These data demonstrate that each MET-type employs distinct connectivity patterns that differ by layer and target cell subclass (Fig 3B).

Of note, we also find that MET-8 cells’ output synapses are significantly smaller than those from MET-4 cells. MET-8 cells form synapses that are 68% the size of MET-4 synapses and 79% of the size of MET-6 synapses (Supplemental Fig 3A).

In Fig 3C, we provide examples of the soma locations of postsynaptic targets (color-coded by the number of synapses received) from each MET-type. This illustrates the differences in the target location and range in number of synapses per connection for each MET-type. MET-8 cells form multi-synaptic connections with L4 and L5 IT pyramidal cells, which can be seen in the cluster of orange-yellow somas directly above the presynaptic cell soma (cyan dot). MET-4 cells target diffusely across most targets but form a few multi-synapse connections in L5. Lastly, MET-6 cells target diffusely across most targets, but form multi-synaptic connections with L5 ET cells located just above the presynaptic cell soma (Fig 3C).

#### IV. Inter-synapse distances of multi-synaptic connections

To determine whether presynaptic Sst-MET-types form spatially clustered synapses onto a postsynaptic target, we quantified the inter-synaptic distances between all synapses of a given pre-post pair. We then built a histogram of those distances across MET-types. We find that most synapses are formed within 150 µm of each other (euclidean distance) (Fig 3D). However, both MET-4 and MET-6 cells have many synapses that are 400 µm apart. These distances may be due to synapses formed onto both the apical and basal dendrites of a target cell or across a wide lateral extent of basal dendrites.

One hypothesis regarding L5 Martinotti cell connectivity is that they form synapses onto both the apical and basal dendrites of L5 excitatory cells (see histograms in Fig 2 and > 400um distances in Fig 3D) to coordinate inhibition across compartments of individual cells. To determine where synapses were formed onto L5 targets, we calculated the percentage of connections from MET-4 and MET-6 with synapses above the middle of L2/3 (presumed onto apical tuft), below the middle of L2/3 (presumed onto basal dendrite), and both above and below the middle of L2/3 (presumed tuft and basal dendrites). We find that most connections that MET-4 cells form with L5 targets (L5 IT: 75%, L5 ET: 80%) are likely onto basal dendrites. Less than 10% of connections are onto apical tufts only and ∼17% of connections for both L5 IT and ET are onto both apical tufts and basal dendrites. For MET-6, most connections are onto basal dendrites only (L5 IT: 77%, L5 ET: 59%) and less than 10% are onto apical tuft only. However, we find approximately 15% of connections onto L5 IT cells span apical tufts and basal dendrites, but ∼34% of connections with L5 ET cells are onto apical and basal dendrites. Thus, the coordinated inhibition onto both the apical and basal dendrites of L5 types by individual Martinotti cells, occurs most commonly between MET-6 and L5 ET cells. However, in no cases observed is it the dominant connectivity motif.

## Limitations

There are a few technical artifacts that will have minor effects on the quantifications of connectivity we report here. First, we estimate that 14% of synapses (see Methods) that were not included in the analysis could be attached to single soma targets with further proofreading. Additionally, the top 10 µm of the cortical surface is not included in these reconstructions due to segmentation errors, however, we find that synapses onto L5 targets peak at a lower depth compared to the peak of synapses onto L2/3 targets. We therefore don’t expect either of these effects to be large enough to change the overall conclusions reported here.

This study supports the use of morphological features from Patch-seq to predict cell type identity for cells in other morpho-containing datasets; however, the sample size of reconstructed EM cells is small and was restricted to the same cortical region (VISp) as the Patch-seq data.

## Discussion

By establishing a morphological feature set aligned across EM and a previously generated, multi-modal Patch-seq dataset, we are able to map cells between the two datasets to predict the connectivity of different types of Sst inhibitory neurons described by their combined morphological, electrophysiological, and transcriptomic properties (MET-types). We can additionally predict the molecular identity and electrophysiological profile of neurons sampled in the large volume EM data. These predictions reveal that MET-types consistently differ in their myelination, synaptic features, and target cell subclass connectivity profiles, which suggests that they play unique roles in the broader cortical circuitry (Fig 4).

**Fig 4:**
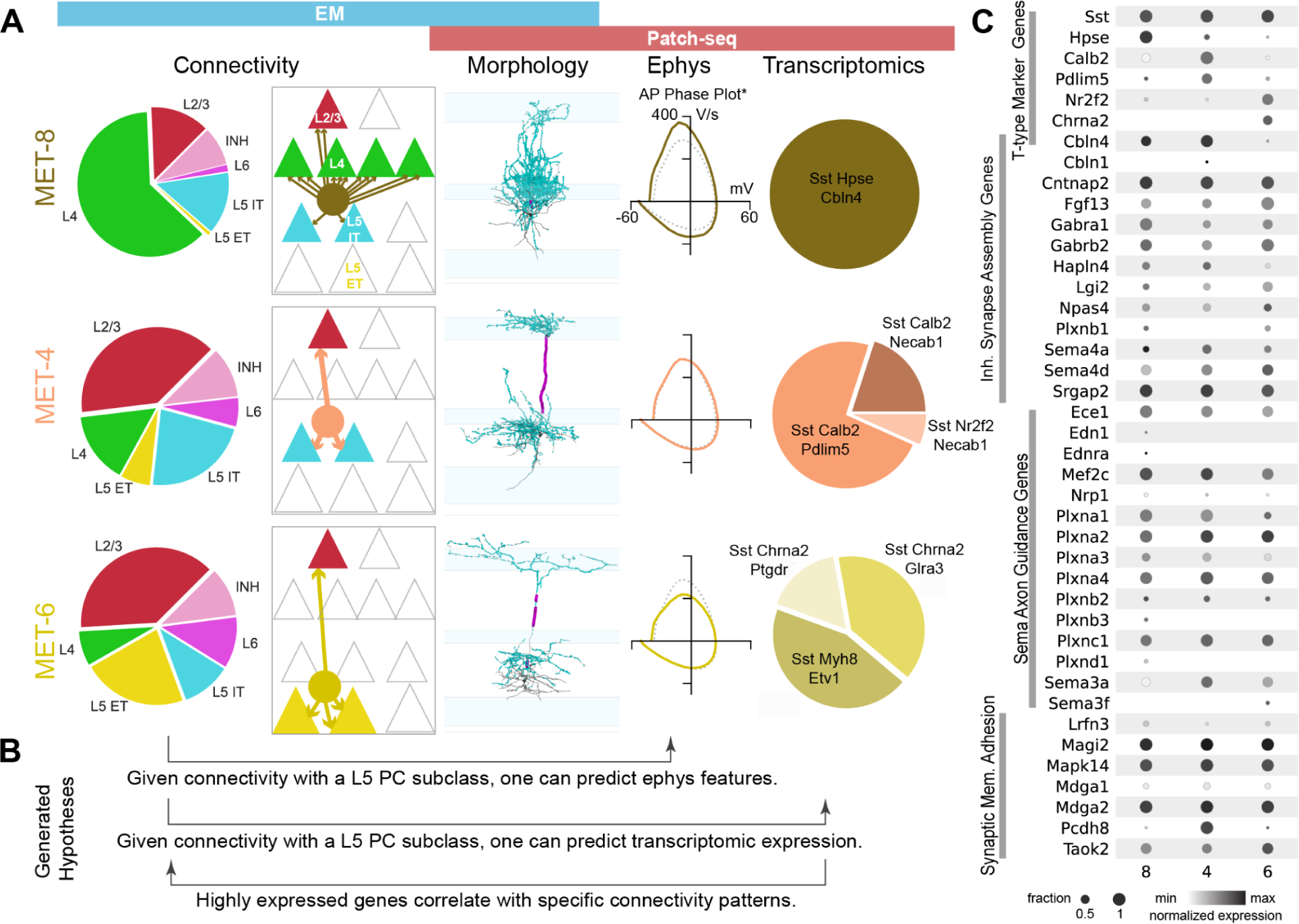
Integrated view of MET-types including modalities from Patch-seq and EM. A. The first column shows the avg. percent of synapses onto each postsynaptic target group. The second column is a schematic summarizing the connectivity motifs observed from the EM data. The third column includes example cells from the EM dataset. The fourth column shows the average AP traces per MET-type (‘data previously published ^21^). The fifth column shows the transcriptomic cell types that comprise previously defined MET-types. B. This integrated view of MET-types now allows us to generate hypotheses such as the role of highly expressed genes in a given transcriptomic type on connectivity patterns. C. Dot-plot showing the proportion (circle size) and expression level (grayscale intensity) of genes used to define the transcriptomic types in MET 8,4,6 as well as genes known to play a role in inhibitory synapse assembly and semaphorin-plexin axon guidance (Gene ontology^58^). Genes such as these might be of interest when testing hypotheses generated from the integrated MET-type view.

### Integrated view of MET-types and connectivity

Previously defined MET-types^21^ have been shown to have unique morphological, electrical, and transcriptomic properties (Fig 4A). When we look across these features, we can now see how MET-8, MET-4 and MET-6 target distinct subclass populations using different connectivity motifs (Fig 4A). We can also generate hypotheses about what features measured in Patch-seq (e.g. electrophysical properties, transcriptomic expression patterns) may correlate with the connectivity patterns observed in EM (Fig 4B). We show expression patterns of a subset of genes known to play a role in various aspects of inhibitory axon guidance and synapse formation ^58^ (Fig 4C). Genes such as these could play a role in setting up and maintaining the distinct morphology and connectivity patterns we observe across these MET-types. Subcellular protein expression and loss-of-function studies during development could be very valuable in determining their role in synaptic patterning.

### Diverse myelination patterns may support distinct inhibitory functions

Though myelination has most frequently been described with respect to long-range projecting excitatory neurons, previous studies have shown that local, inhibitory neurons ^45, 59^ including Martinotti cells^47^, can be myelinated. Here we characterize myelination patterns across multiple, distinct Sst-MET-types and find significant differences in the number and total length of myelinated segments (Supplementary Fig. 2C). Importantly, reduced myelination onto a different inhibitory cell subclass (Pvalb+) has been shown to reduce firing rates and conduction velocity ^45, 60^. It seems likely that the myelination of putative Sst+ cells may similarly influence their firing properties. Given this observation, the increased myelination of the main ascending axon stalk of MET-4 cells may help synchronize inhibition of L2/3 and L5 IT pyramidal cell targets and/or increase the speed of inhibition onto L2/3 targets of MET-4 versus MET-6 cells (Fig 2A, Supp Fig 3B). The observed difference in myelination across these MET-types suggests myelination may vary in a cell type specific manner and might provide another marker of cell type identity in EM datasets.

### Inhibitory output synapse size may support distinct circuit roles

We find that MET-4 cells have significantly larger output synapses and MET-8 cells form significantly smaller output synapses than the other two MET-types (Supplemental Fig 3A). Correlated slice electrophysiology and EM of synaptically-connected excitatory cells has shown a linear relationship between chemical synapse size and synaptic strength ^61^. It might be possible that an individual MET-8 synapse is weaker than any individual MET-4 or MET-6 synapse. However, we also find that MET-8 cells form significantly more output synapses than either MET-4 or MET-6 cells and so a single synapse analysis might be under-counting their inhibitory influence on the circuit.

MCs can form multiple synapses onto a single target, so we calculated a cell’s average connection size (the sum of synapse sizes onto each postsynaptic target divided by the number of postsynaptic targets). We find that MET-8 cell connections are larger than those of MET-6 (though not MET-4). It is possible that the greater number of synapses onto individual targets may be related in part to the smaller synapse size. This pattern of connectivity (many small synapses per target) may reflect a ‘hard wired’ program or could be an outcome of homeostatic plasticity.

Inhibitory synapses onto excitatory cells in cortical and hippocampal circuits scale homeostatically in response to changes in activity at the circuit ^62–64^ and cell level ^65–68^ though this has mainly been described for the Pvalb+ cell population ^66, 67^. The number and size of inhibitory synapses onto a target may specifically balance the excitation impinging onto the same cell and shape the postsynaptic cell response. In chicken auditory nucleus, inhibition varies along the tonotopic axis, which is important for shaping the timing and dynamic range of postsynaptic responses^68^. Thus, inhibitory synapse size may be a function of homeostatic plasticity and/or be suited to the features encoded by the postsynaptic targets of each MET-type.

#### “Blanket of inhibition” in some, but not all layers

Previous studies found dense connectivity from L2/3 *Sst+* cells onto L2/3 pyramidal cell targets creating a “blanket of inhibition” in the cortex^33^ and predicted “non-specific” connectivity onto most cell types^14^. We examined this possible connectivity in L4 and L5. We find that some predicted *Sst+* types very densely target cells with nearby somas (MET-8 targets nearly all nearby L4 pyramidal cells); however, we also see that other MET-types differentially target cells within their resident layer. We therefore observe a “blanket of inhibition” from the MET-8 cells onto L4 cells, but not onto other target subclasses of MET-8 or from other MET-types. While a “blanket of inhibition” is visible when considering specific pre and postsynaptic targets, this connectivity pattern is not the default for all predicted *Sst+* cells across all targets.

#### Observed complementary networks of inhibition (L4 vs L5)

*Sst+* cells in L4 and 5 with distinct axonal morphology have distinct intrinsic physiology ^21, 69^ and even opposite behavior during whisking^37^. Some studies found that Sst cells specifically innervate excitatory targets in one layer but not the other (*Sst*+ cells innervating L4 will target L4 and not L5 pyramidal cells and vice versa)^32, 38, 39^. Whereas other studies found that a subset of L4-innervating Sst cells target inhibitory rather than excitatory cells ^37, 70^.

We find EM *Sst+* MET 4,6 & 8-types predominantly contact excitatory and not inhibitory targets. However, predominantly inhibitory targeting *Sst*-types may be present in the data but not yet identified, and it is possible that many cells collapsed into the “inhibitory” category may differ across MET-types. A concurrent paper^40^ finds that perisomatic-targeting inhibitory cells (likely basket/fast spiking cells) are the major inhibitory target of L4 dendrite targeting cells (likely *Sst+*). We find cells (MET 4 & 6) that target L5 but not L4 pyramidal cells and cells (MET-8) that inhibit L4 and not L5 pyramidal cells, supporting the previous descriptions of parallel complementary inhibitory networks across layers 4 and 5^38^.

#### MET-type connectivity aligns with previous physiological findings and connectivity motifs

Hpse Cbln4 cells have been previously reported in mouse primary somatosensory and visual cortex and were shown to connect with L4 and not L5 pyramidal cells ^29, 38^. Scala et al.,^29^ also observed morphological differences between Hpse Cbln4 neurons in somatosensory (no L1 projection) versus visual cortex (some L1 projection). If MET-8 cells are Hpse Cbln4+^21^ these EM data recapitulate both the morphological and connectivity findings in mouse VISp.

Recent work in mouse VISp finds that bulk optogenetic activation of virally labeled Sst Calb2 cells (many of which would likely be classified as Sst-MET-4 cells) produce much larger amplitude iPSCs onto L5 ET than IT cells^39^, whereas we find that the MET-4 group forms more output synapses onto L5 IT cells and has similar connection probabilities to L5 IT and ET cells. The differences in our findings could be accounted for by several key differences in approach: the Sst Calb2 population likely includes cells from both MET-3 and MET-4 groups, whereas we focused on MET-4; their functional measures of inhibition are made postsynaptically and reflect convergence of multiple presynaptic Sst cells, whereas we measure the number of synapses from individual presynaptic cells; and lastly their postsynaptic cell subclasses are defined by retrograde labeling and may reflect more restricted populations (e.g., retrosplenial cortex targeting neurons), whereas we used local somatic features to define the postsynaptic subclass. Alternatively, this could reflect that MET-4/Sst Calb2 cells have functionally stronger synapses with ET cells than IT cells per individual synapse potentially due to differences in presynaptic release probability, receptor composition, receptor density, or dendritic integration. Future targeted studies to examine the functional dynamics of individual synapses onto each excitatory target population would help to resolve this.

Previous studies have also found that *Chrna2*+ Martinotti cells in L5 of auditory cortex preferentially inhibit Type A (thick-tufted/L5 ET cells) but not Type B (thin-tufted/L5/6 IT cells) cells^36^ and optogenetic mapping of Sst-Myh8 cells (genetically targeted using Chrna2-cre) shows that they more strongly inhibit L5 ET cells than IT cells ^39^. We find MET-6 cells, which contain *Chrna2*+ cells ^21^, form more synapses onto L5 ET than IT cells. Thus, the MET-6 type connectivity aligns with that of *Chrna2*+ Martinotti cells^36^.

These findings suggest that cells which exhibit similar biases in connectivity across sensory cortical regions (visual, somatosensory, auditory) may also share molecular features (Hpse Cbln4, *Chrna2*+) and may even be the same cell-type. Transcriptomic profiling in the mouse brain reveals that inhibitory cell types are largely conserved across isocortical regions^8^ and hippocampal formation (e.g. Sst Etv1 cells are seen in both isocortex and hippocampal formation)^71^. Further work is needed to compare output connectivity patterns of molecularly identified Sst+ neurons across cortical regions.

Frequency-dependent di-synaptic inhibition (FDDI) is a well-described functional circuit motif characterized by a single L5 pyramidal cell (specifically thick-tufted/ET) exciting L5 Martinotti cells, which, in turn, inhibit surrounding L5 pyramidal cells (typically thick-tufted/ET)^34, 35, 72–74^. FDDI has also been shown to occur when aL2/3 pyramidal cell synapsing onto L5 Martinotti cells inhibits surrounding L2/3 pyramidal cells^75^. MET-6 type cells synapse onto both L2/3 and L5 ET cells and therefore may participate in FDDI.

Finally, we have demonstrated that local morphological features from mouse primary visual cortex can be used to link cell types across datasets. Linking these cell type identities enables the investigation of synaptic connectivity with respect to morphology, electrophysiology, and transcriptomic expression. As larger serial-section electron microscopy datasets are generated, this approach can be extended to cell types in different brain regions. Measuring the synaptic connectivity of identified cell types will facilitate future work aimed at characterizing the behavior of these cell types within local circuits and across the brain.

## Methods

### EM dataset generation and image alignment ^30^

All animal procedures were approved by the Institutional Animal Care and Use Committee at the Allen Institute for Brain Science or Baylor College of Medicine. In brief, a large-scale serial-section electron microscopy dataset was collected and imaged using automated transmission electron microscopes^76^. The data above is from a sub-volume representing 65% of the original EM volume with images of ∼4x4x40 nm/pixel resolution. These images were segmented into meshes using convolutional neural networks and subsequent agglomeration^77^. The EM images and meshes are visualized in Neuroglancer. These meshes can be proofread (merged/split) within the ChunkedGraph system ^40, 78^ Neuroglancer framework to facilitate proofreading of cells.

### Correcting and generating representations of cells

Meshes underwent skeletonization (skeleton originated from a defined soma point) to generate a list of branch and end points for each mesh, visible in Neuroglancer^40^. Each branch point was manually inspected. True branch points were left alone and false branch points (often due to overlapping processes from distinct cells) were split using Neuroglancer tools. Subsequently, each endpoint was manually inspected. True endpoints were left alone and false endpoints (premature end of a process) were extended by an expert annotator would follow the process along the EM imagery to a natural ending (bouton, tapered end) or until the process could no longer be reliably extended (e.g. edge of block).

### Morphological analysis and MET-type prediction

Soma position, pia, white matter, and laminar borders were manually drawn. For Patch-seq cells a 20X brightfield and fluorescent image of DAPI (4′,6-diamidino-2-phenylindole) stained tissue was used ^21^ and for EM cells drawings were made in Neuroglancer on a single EM z-plane that contained the soma of the cell of interest. These polygons were then exported to be used for feature calculation.

Morphological features were calculated as previously described^16^, using features that were derived from prior studies ^15, 79^. Features were calculated using the skeleton keys python package (https://github.com/AllenInstitute/skeleton_keys). Features were extracted from neurons aligned in the direction perpendicular to pia and white matter. Laminar axon histograms (bin size of 5 microns) and earth movers’ distance features require a layer-aligned version of the morphology where node depths are registered to an average laminar depth template.

A random forest classifier was trained on the morphological features of multiple inhibitory cell types from a previously published Patch-seq dataset^21^ with MET-type labels. For 500 iterations a random subsample (95%) of the Patch-seq data was selected with probabilities according to MET-type class size (a Patch-seq cell from a well represented met-type was more likely to be omitted). MET-types with 5 or fewer specimens were exempt from subsampling. In each iteration, a unique random forest classifier was fit with subsampled Patch-seq data and MET labels were predicted for EM cells. Out of bag scores were recorded for each iteration (Mean, Stdev = 0.58, 0.013). The final MET assignment was given as the most frequently predicted MET label for each cell (Supplementary Table 1). We used these predicted MET-type labels (if predicted into that MET-type >55% of the time) to group cells for subsequent analysis.

### Identifying synapses and postsynaptic targets ^43, 50^

Synapses and their pre- and post-synaptic meshes in the EM dataset were previously algorithmically detected ^50^. These data also included the automatically detected synapse size (number of voxels per synapse)^50^.

Cell subclass identities were assigned to all meshes with single somas (i.e. individual cells) in the EM dataset using a svm classifier trained somatic and nuclear features ^43^. This classifier was then applied across the EM dataset to generate predicted cell-type identities for most cells. We use these identities in all plots shown above, however, we manually inspected 5% of all output synapses per cell to confirm the presence of true synapses and to determine the postsynaptic target cell identity. There was broad agreement between the automated and manual cell-typing except for a specific disagreement of L2/3 vs L4 identity for targets of the predicted MET-8 cell type due to differences in layer boundaries used by manual vs automated methods (Supplementary Table 3).

The large majority of synapses from individual cells were onto postsynaptic targets which contained single somas in the reconstruction (Supplemental Table 2). Of the synapses which were not onto single soma targets, estimated less than 15% are onto multi-soma targets, and 14% are onto orphan segments which are not presently connected to a soma and ∼4% of synapses are onto targets with somas outside of the block. We spot checked ∼190 orphan segments spread across at least 3 reconstructions per MET-type, and attempted to proofread them to see if it was possible to connect it to a soma. The true distribution of synapses and rates of connectivity onto excitatory cells might be marginally higher than we measured in the automated analysis. Further work on cell type characterization and proofreading will improve these numbers in the future.

### Quantification of myelination

Using automatically detected synapses, annotators visualized all output synapses on a given presynaptic cell in Neuroglancer. Regions lacking synapses were manually inspected in the EM imagery. If myelination was seen, an annotator marked the start and end point of each myelinated segment in Neuroglancer to generate a line. The number of these annotations was summed to determine the number of myelinated segments per cell. The length of each annotation was summed to determine the distance of myelinated axon per cell.

### Apical Tuft vs Basal dendrites connectivity

Taking the previously drawn cortical layers for each reconstructed cell, we calculated the average depth for the middle of L2/3 (average of upper and lower boundaries of L2/3). We used this depth to calculate the % of connections (for each pre-post pair - only considering MET-4 and MET-6 cells) with synapses that were all above (presumed tuft only), all below (presumed basal only), or spanned the middle of L2/3 (presumed apical tuft and basal synapses). These percentages were averaged for each MET-type and reported.

### Statistics

Comparisons across multiple MET-types were performed using non-parametic Krusall-Wallis tests followed by Conover post-hoc tests with Bonferonni corrections for pairwise comparisons. P values are reported for both the Krusall-Wallis and posthoc tests. Errors reported are standard error of the mean (s.e.m) unless otherwise indicated.

## Data and code Availability

Analysis for this paper was performed on version 500 of the dataset, a snapshot taken on September 19th, 2022 at 8:10am UTC. The mm^3^ EM dataset is publicly available at https://www.microns-explorer.org/cortical-mm3. Analysis code will be made publicly available on Allen Institute Github repository (forthcoming). Analysis was performed using Python 3.x and made extensive use of the following packages and libraries: CAVEclient (https://github.com/seung-lab/CAVEclient), CloudVolume ^80^, MeshParty^81^, skeleton_keys (https://github.com/AllenInstitute/skeleton_keys) to extract and analyze morphological features. We used the following libraries for visualization and analysis:Matplotlib^82^, Seaborn^83^, Numpy^84^, Pandas^85^, VTK^86^, Scipy^87^, Scikit-posthocs^88^, Scikit-learn^89^.

## Acknowledgements

The work was supported by the Intelligence Advanced Research Projects Activity (IARPA) via Department of Interior/ Interior Business Center (DoI/IBC) contract numbers D16PC00003, D16PC00004, and D16PC0005. The U.S. Government is authorized to reproduce and distribute reprints for Governmental purposes notwithstanding any copyright annotation thereon. NMdC, FC and RCR also acknowledge support from NIH RF1MH125932 and from NSF NeuroNex 2 award 2014862.

The research was also supported by several grant awards from institutes under the National Institutes of Health (NIH), including award numbers R01EY023173 from The National Eye Institute and U01MH105982 from the National Institute of Mental Health and Eunice Kennedy Shriver National Institute of Child Health and Human Development to H.Z. The content is solely the responsibility of the authors and does not necessarily represent the official views of NIH and its subsidiary institutes. This work was funded by the Allen Institute for Brain Science. We thank Allan Jones and Rui Costa for leadership and guidance and dedicate this paper to the vision, encouragement, and long-term support of our founder, Paul G. Allen.

Special thanks to Ashwin Bhandiwad, Yoni Browning, Thomas Chartrand, Travis Hage, Jeremy Miller, Jenna Schardt and Stephanie Seeman for their constructive feedback and conversations.

## Author Contributions

C.R.G and S.A.S. conceived the project. C.M. S-M, L.E., A.B, D.B, J.B., D.B., D.K., S.K., G.M., M.T., R.T., W.Y., J.A.B., M.A.C., S.D., A.H., Z.J., C.J., N.K., K. Lee, K. Li, R.L. T.M., E.M., S.S.M., S.M., B.N., S.P., W.S., N.L.T., W.W., J.W., S.Y., H.S.S., R.C.R., F.C., and N.M.dC. contributed to the design, collection, alignment, segmentation, and development of software to analyze the electron microscopy dataset. J.B., N.G., T.J., B.L., P.R.N, S.A.S. and H.Z. contributed to the design and J.B., N.G., T.J., B.L., S.A.S. and H.Z. contributed to the leadership on the generation of the Patch-seq dataset. C.R.G., C.M. S-M., R.D., G.W., and A.B. contributed to data curation. A.M. and S.S. contributed code for analysis. C.R.G., M.M., N.G., and F.C. contributed to data analysis. C.R.G. wrote the manuscript. C.M. S-M., R.D., T.J., H.Z., F.C., N.M.dC., and S.A.S. revised the manuscript.

## Corresponding Authors

Correspondence to Forrest Collman, Nuno da Costa, and Staci Sorensen.

---

**Supp. Fig. 1.**
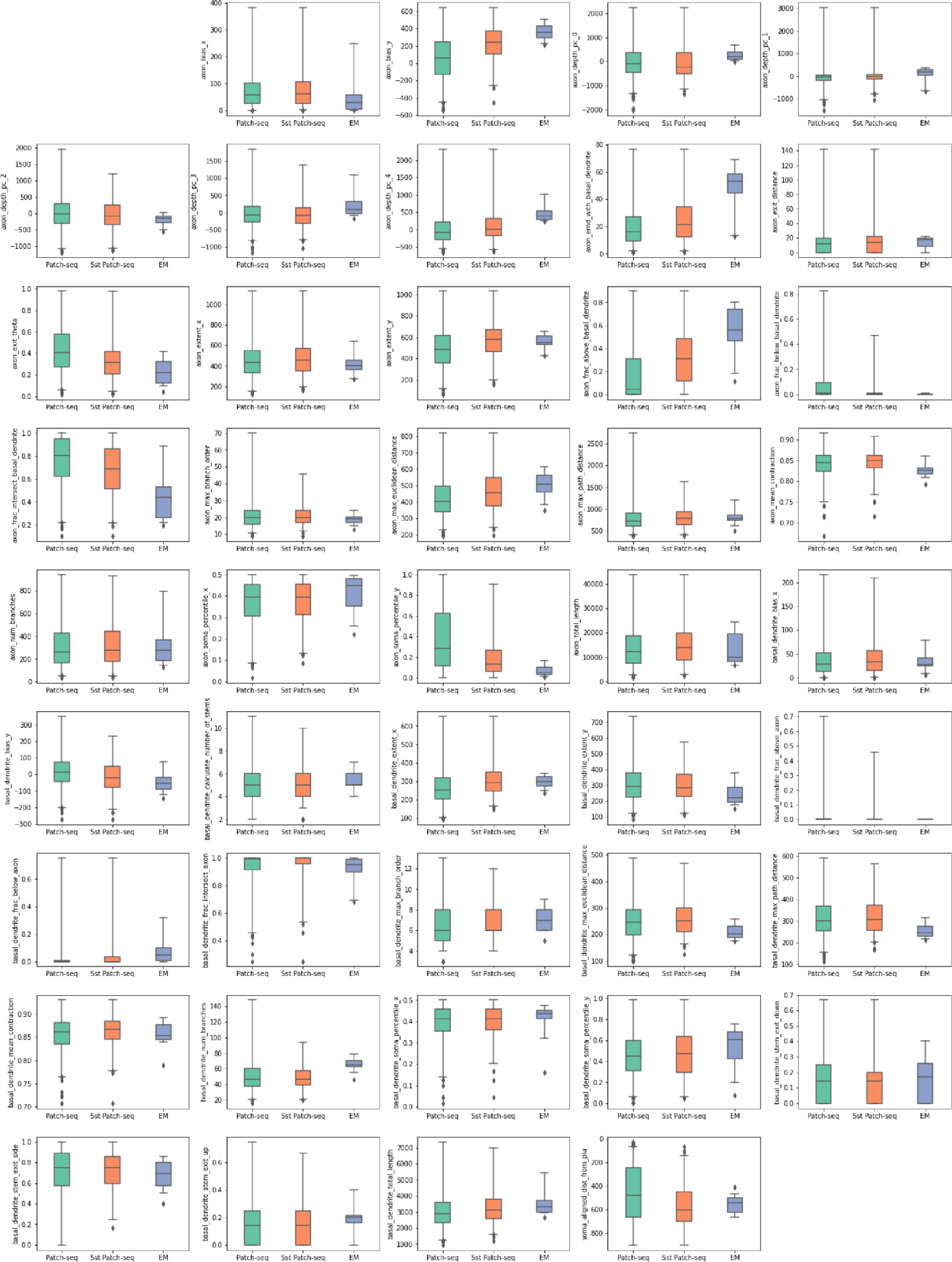
Box-whisker plots (whiskers= range) comparing the morphological features of all inhibitory Patch-seq vs just the Sst+ Patchseq vs EM cells.

**Supp. Fig. 2.**
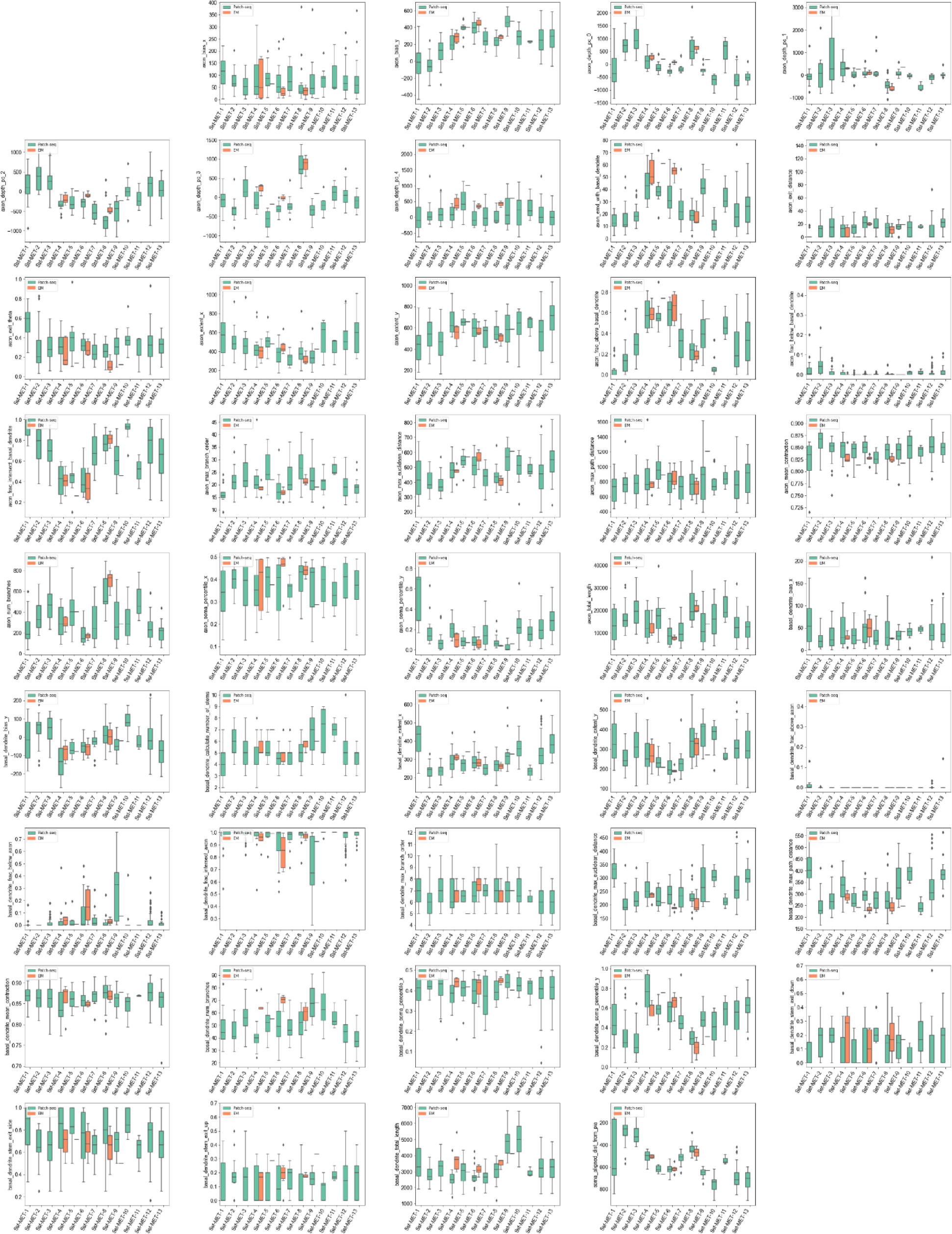
Box-whisker plots (whiskers = range) comparing the morphological features of Sst+ Patchseq vs EM cells by MET or predicted MET-type.

**Supp. Fig. 3.**
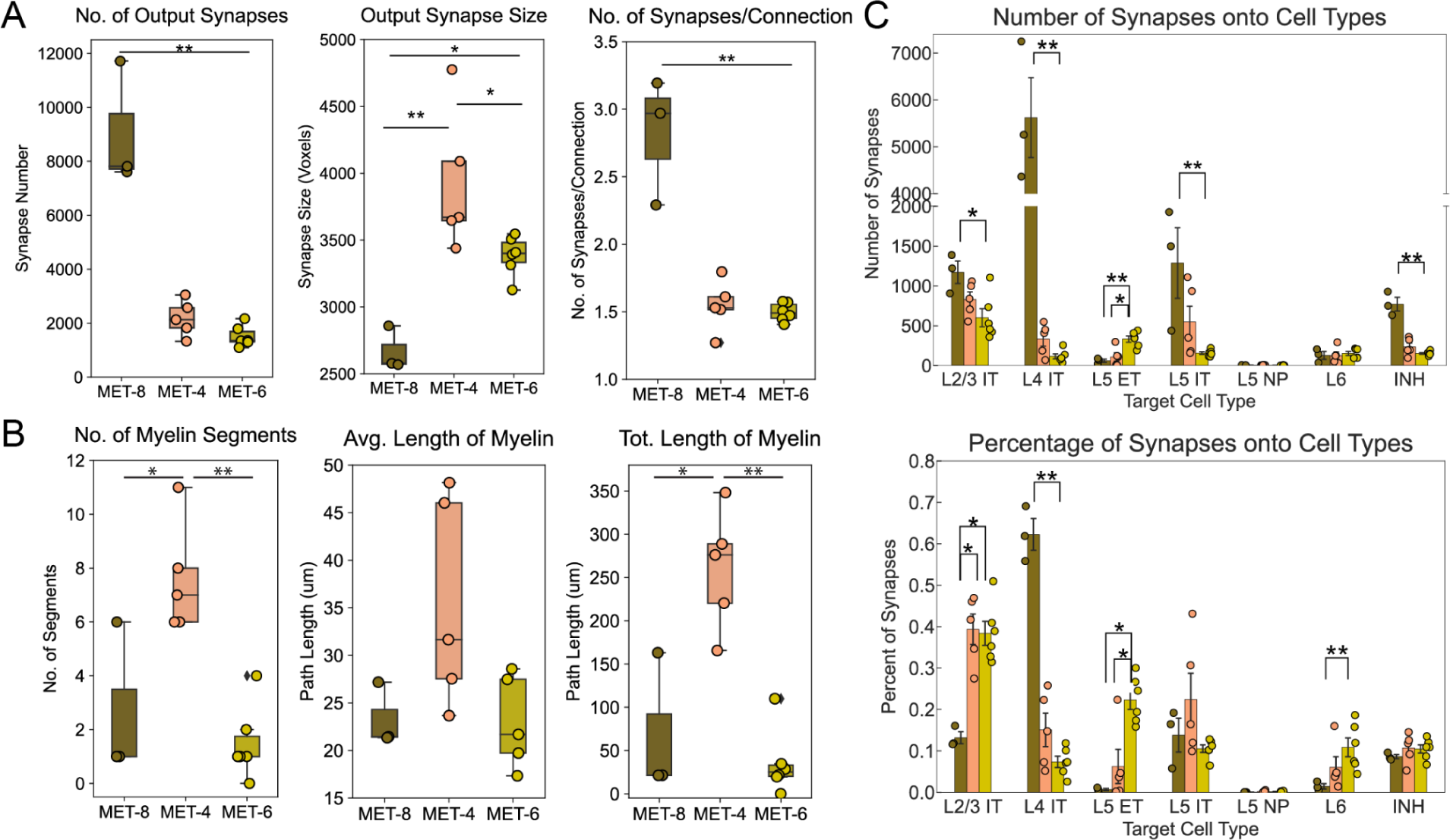
Quantification of output synapse and myelination features by MET-type. A. Quantification of number of output synapses across predicted MET-types [Kruskal-Wallis test: p = 0.02; Conover (Bonferroni correction) 8 v 6: p = 0.003], size [Kruskall-Wallis test: p = 0.006, Conover (Bonferroni correction) 8 vs 4 p = 0.0002, 8 vs 6 p = 0.025, 4 vs 6 p = 0.011], number of synapses per connection [Kruskall-Wallis test: p =0.028, Conover (Bonferroni correction) 8 vs 6: p = 0.01]. B. Quantification of number [Kruskal-Wallis: p=0.012; Conover test [Bonferroni correction]; 8 vs 4: p = 0.026; 4 vs 6: p=0.002)], average length, and total length [Kruskal-Wallis: pvalue=0.011; Conover test [Bonferroni correction] 8 v 4: p = 0.012; 4 v 6: p = 0.002] of myelinated segments across predicted MET-types. C. Quantification of the total number and percentage of output synapses each target cell type. Bars indicate significant differences (p < 0.05 Krusall-Wallis for group and Conover post-hoc, * p<0.05, ** p<0.01)

**Supp. Fig 4.**
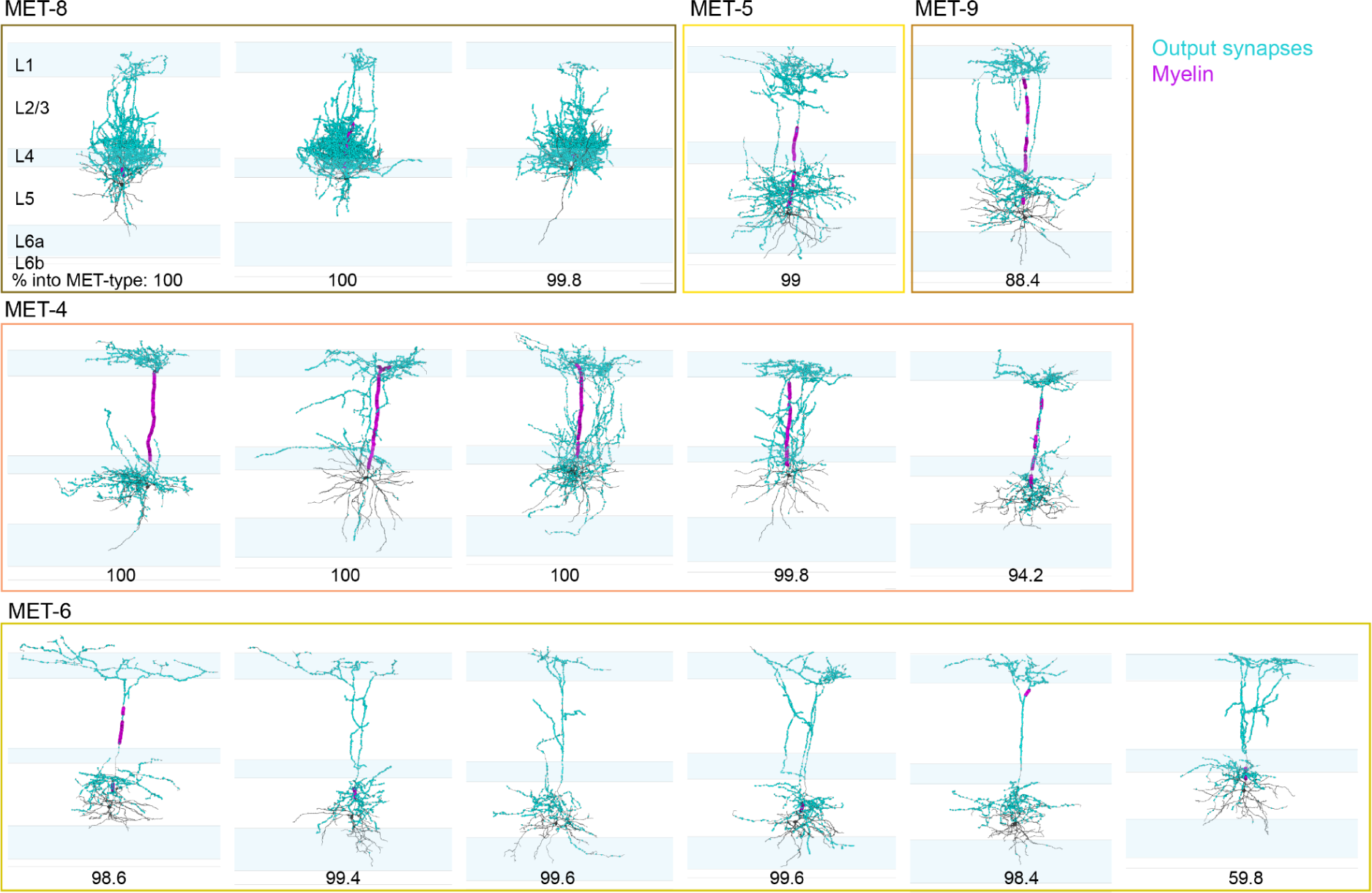
Output synapses and myelination patterns of all EM MCs grouped by predicted MET-type. Percentage (out of 500 runs) that the cell was predicted into that MET-type displayed below.

**Supp. Fig 5.**
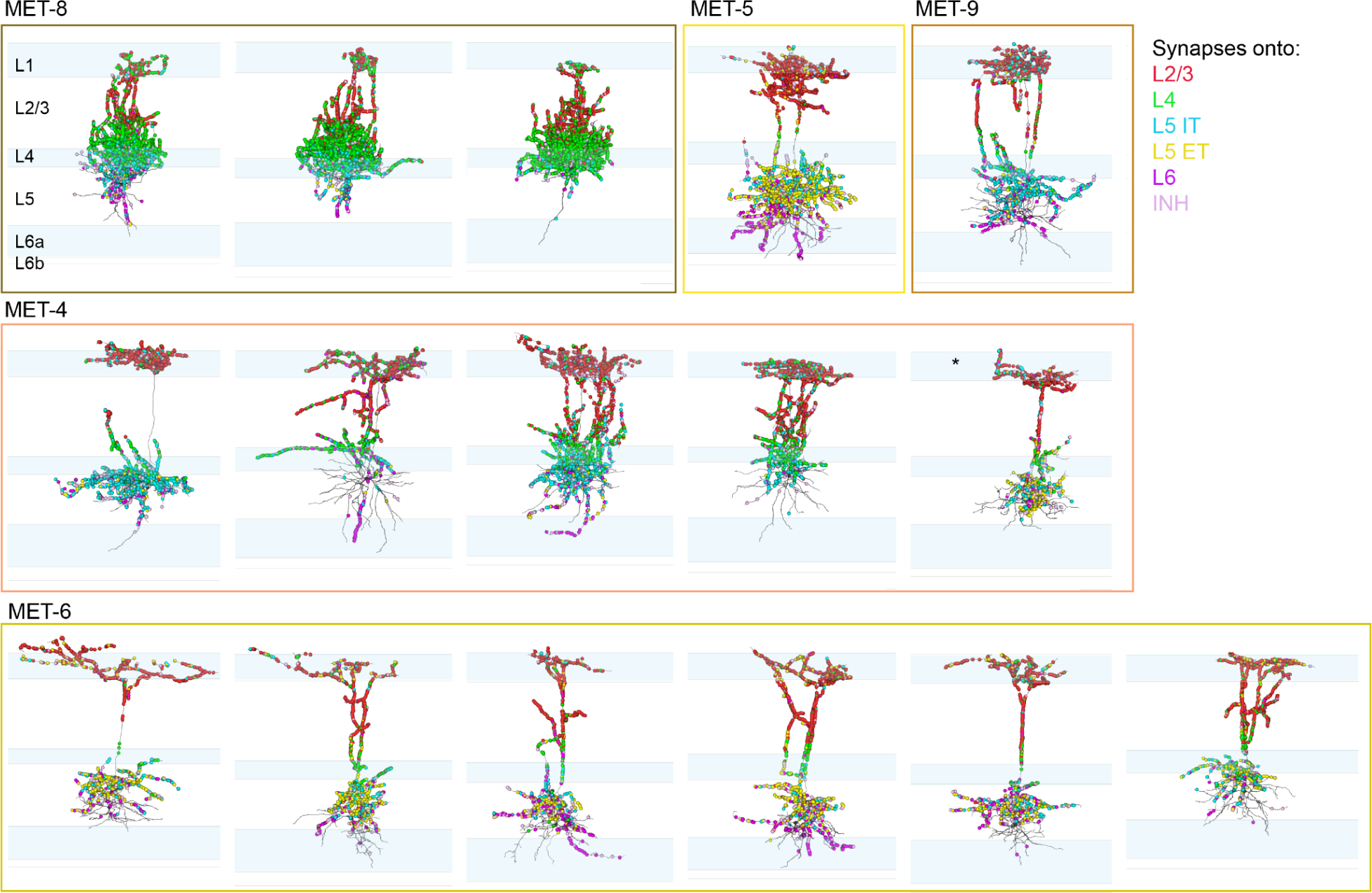
Output synapses color-coded by postsynaptic cell subclass of all EM MCs grouped by predicted MET-type. * denotes cell with dominant L5 ET targeting in contrast to the rest of the MET-4 cells.

**1.** Table showing cell type id, predicted MET-label, the fraction (out of 500 runs) the cell was predicted to that MET-type.

| Cell ID | Predicted MET-type | Fraction |
| --- | --- | --- |
| 864691135013417622 | Sst-MET-4 | 1 |
| 864691135467660940 | Sst-MET-4 | 1 |
| 864691135699487522 | Sst-MET-8 | 1 |
| 864691135925834510 | Sst-MET-8 | 1 |
| 864691136118647832 | Sst-MET-4 | 1 |
| 864691135544588584 | Sst-MET-8 | 0.998 |
| 864691135577202181 | Sst-MET-4 | 0.998 |
| 864691135374222153 | Sst-MET-6 | 0.996 |
| 864691136116457636 | Sst-MET-6 | 0.996 |
| 864691135058985115 | Sst-MET-6 | 0.994 |
| 864691135754152141 | Sst-MET-5 | 0.99 |
| 864691135988665856 | Sst-MET-6 | 0.986 |
| 864691136618564493 | Sst-MET-6 | 0.984 |
| 864691135785592004 | Sst-MET-4 | 0.942 |
| 864691135341516741 | Sst-MET-9 | 0.884 |
| 864691135118298333 | Sst-MET-6 | 0.598 |

**2.**
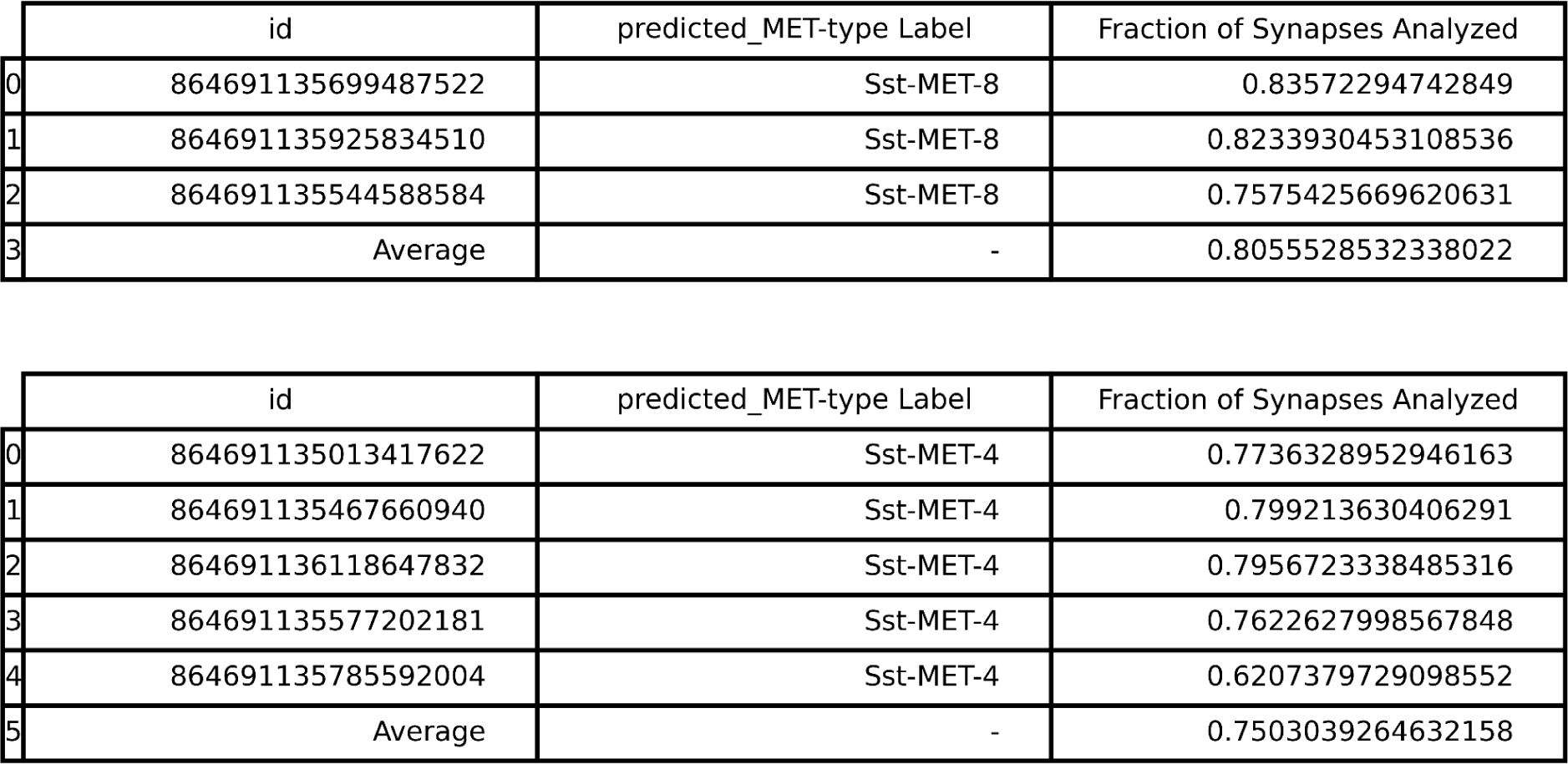
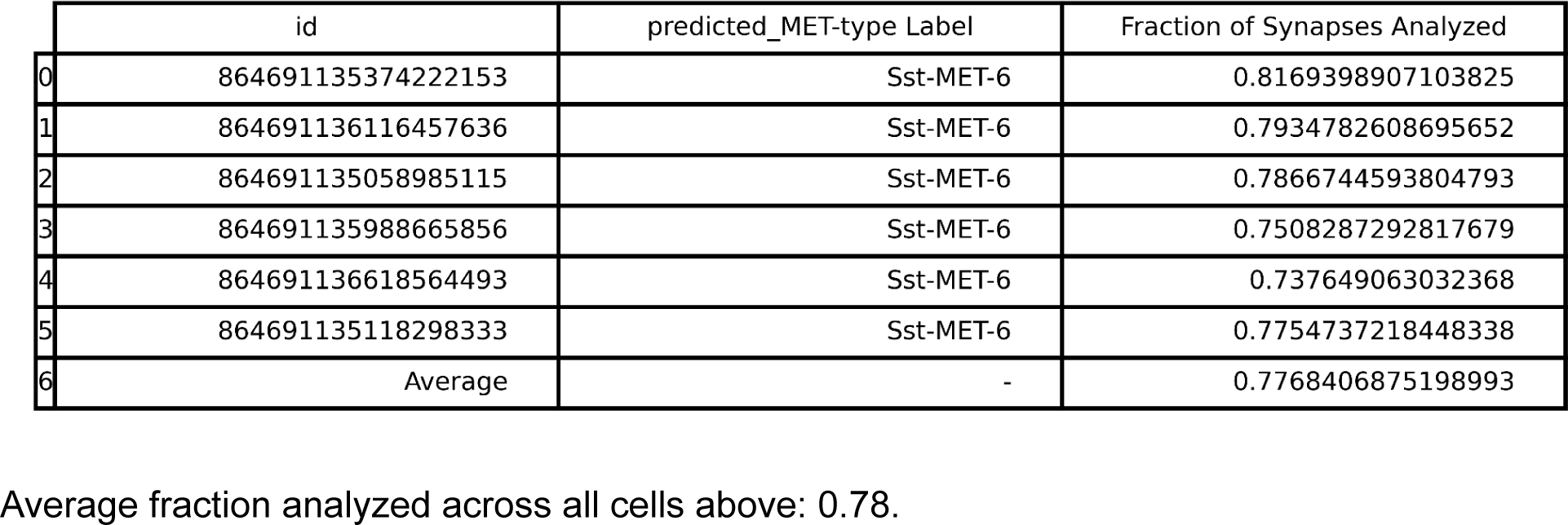
Percentage of output synapses that were onto single-soma objects with cell-type predictions (Low numbers may be due to cell location relative to tissue boundaries)

|  | id | predicted_MET-type Label | Fraction of Synapses Analyzed |
| --- | --- | --- | --- |
| 0 | 864691135699487522 | Sst-MET-8 | 0.83572294742849 |
| 1 | 864691135925834510 | Sst-MET-8 | 0.8233930453108536 |
| 2 | 864691135544588584 | Sst-MET-8 | 0.7575425669620631 |
| 3 | Average | - | 0.8055528532338022 |
| 0 | 864691135013417622 | Sst-MET-4 | 0.7736328952946163 |
| 1 | 864691135467660940 | Sst-MET-4 | 0.799213630406291 |
| 2 | 864691136118647832 | Sst-MET-4 | 0.7956723338485316 |
| 3 | 864691135577202181 | Sst-MET-4 | 0.7622627998567848 |
| 4 | 864691135785592004 | Sst-MET-4 | 0.6207379729098552 |
| 5 | Average | - | 0.7503039264632158 |
| 0 | 864691135374222153 | Sst-MET-6 | 0.8169398907103825 |
| 1 | 864691136116457636 | Sst-MET-6 | 0.7934782608695652 |
| 2 | 864691135058985115 | Sst-MET-6 | 0.7866744593804793 |
| 3 | 864691135988665856 | Sst-MET-6 | 0.7508287292817679 |
| 4 | 864691136618564493 | Sst-MET-6 | 0.737649063032368 |
| 5 | 864691135118298333 | Sst-MET-6 | 0.7754737218448338 |
| 6 | Average | - | 0.7768406875198993 |

Average fraction analyzed across all cells above: 0.78.

**3.**
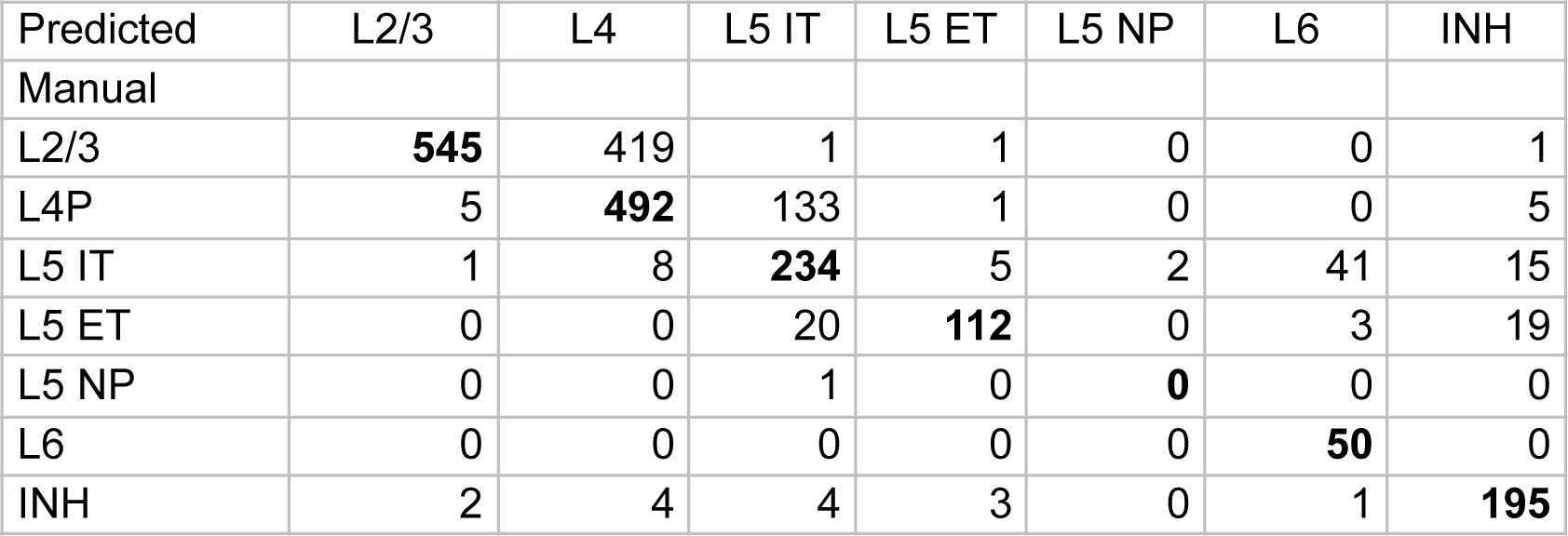
Table showing differences between manual (expert annotator) vs predicted subclass calls from Elabbady et al.^43^. The largest disagreement results from the human annotator referencing layer drawings from different z-planes throughout the dataset which features a larger L2/3 and more restricted L4, whereas the classifier used labels derived from a cortical column which resulted in more cells being called L2/3 that the classifier predicted at L4. Diagonal elements are bolded.

| Predicted | L2/3 | L4 | L5 IT | L5 ET | L5 NP | L6 | INH |
| --- | --- | --- | --- | --- | --- | --- | --- |
| Manual |  |  |  |  |  |  |  |
| L2/3 | **545** | 419 | 1 | 1 | 0 | 0 | 1 |
| L4P | 5 | **492** | 133 | 1 | 0 | 0 | 5 |
| L5 IT | 1 | 8 | **234** | 5 | 2 | 41 | 15 |
| L5 ET | 0 | 0 | 20 | **112** | 0 | 3 | 19 |
| L5 NP | 0 | 0 | 1 | 0 | **0** | 0 | 0 |
| L6 | 0 | 0 | 0 | 0 | 0 | **50** | 0 |
| INH | 2 | 4 | 4 | 3 | 0 | 1 | **195** |

